# Effects of the copper IUD on composition of the vaginal microbiota in the olive baboon

**DOI:** 10.1101/720185

**Authors:** AJ Eastman, D Sack, D Chai, CM Bassis, KA Carter, VB Young, IL Bergin, JD Bell

## Abstract

**Objectives:** Assess the impact of transcervical insertion, use, and removal of copper intrauterine devices (Cu-IUD) on baboon physiology (*e.g*. weight, menstruation) and vaginal microbiota.

**Study design:** Vaginal swabs were taken before insertion (pre-IUD), during IUD use (IUD), and after removal (post-IUD) and microbiota assessed by 16S rRNA-encoding gene sequence analysis.

**Results:** No animals showed physical changes or discomfort during pre-IUD, IUD, or post-IUD phases. There were no changes to the microbiome associated with insertion or use of Cu-IUD over 16 weeks, although removal resulted in perturbation to community structure.

**Conclusions:** Baboons tolerate Cu-IUD insertion with minimal device-associated changes to their vaginal microbiome throughout use and have no significant changes to their physiology or menstrual cycle during any phase.

**Implications:** A baboon model of Cu-IUD may allow investigations into the intersection of Cu-IUDs, reproductive tract disorders and pathogens that would not be possible in human studies.

## 1. Introduction

A non-human primate (NHP) model for copper intrauterine devices (Cu-IUD) that mimics all aspects of human Cu-IUD usage, including both the device and transcervical insertion, has been elusive. Macaques have a tortuous cervix, which prevents transcervical IUD insertion, and their small size may make device tolerance challenging. The olive baboon (*Papio anubis*) is larger and has a straight cervical canal, making it an ideal animal model for intrauterine contraception. While our lab previously developed a baboon model for hormonal IUDs [1], the size of the Cu-IUD insertion device and the different composition and mechanism of action of the Cu-IUD requires careful characterization. This study was designed to determine 1) whether transcervical insertion of the Cu-IUD is possible without damaging the animal; 2) physiologic impact on the primate due to insertion, use, and removal of the device including weight, temperature, respiration, impact on menstrual cycle; 3) expulsion of the device due to unforeseen factors; and 4) potential changes to the vaginal microbiome due to insertion, use, or removal of the device.

*Lactobacillus* sp. often dominate human vaginal microbiota and are considered healthy and which facilitate low pH and high lactic acid levels thought to prevent outgrowth of other microbes. In humans, the Cu-IUD is frequently associated with– but not proven to cause– an increase in bacterial vaginosis [2], which could suggest that the Cu-IUD alters the vaginal microbiome to become less stable. In contrast, the normal baboon vaginal microbiota is diverse and reminiscent of a BV state, although they do not show symptoms of BV such as discharge or odor [3]. A NHP model for Cu-IUDs would permit studies regarding the impact of Cu-IUDs on the biology of the female reproductive tract [4] and the relationship between intrauterine contraception and infectious agents (*e.g. Chlamydia trachomatis*, HIV) that may be ethically impossible in humans.

Our laboratory previously developed a NHP model of levonorgestrel-releasing intrauterine system (LNG-IUS) with olive baboons, which assessed changes to the vaginal microbiome in baboons with LNG-IUS. We found inter-individual differences, but no consistent device-associated shift in microbial communities [5]. Animals with LNG-IUSs had a moderate shift towards more stable microbial communities over time relative to their baseline pre-LNG-IUS community variation [5]. In this study, we assessed the impact of transcervical insertion, use, and removal of copper intrauterine devices (Cu-IUD) on baboon physiology (*e.g*. weight, menstruation) and analyzed abundance and diversity of the vaginal microbiota over 24 weeks.

## 2. Methods

### 2.1 Regulatory approval

The Institutional Review Committee at the Institute of Primate Research (IPR) in Nairobi, Kenya (NIH Office of Laboratory Animal Welfare foreign assurance A5796) approved this study. This study received an off-site exemption from the University of Michigan University Institutional Animal Care and Use Committee. An export permit was obtained from the Kenyan Wildlife Service and import permits were obtained from the US CDC [5].

### 2.2 Study site, population, design, sampling

This study was conducted at the IPR (for IPR description, see [5]). Five wild-caught sexually mature female olive baboons (*P. anubis*) were utilized in this study, quarantined, examined, housed individually, and vaginally sampled as previously described [5]. The baboons were between 10.5 and 16 kg at the beginning of the study. IUDs were inserted based on manufacturer instructions with the modification that cervical length was assessed as at least 5 cm consistent with our previous model of baboon transcervical IUD insertion. During insertion, betadine was applied exclusively to the cervix to prevent contamination of the upper reproductive tract as analogous to human insertion. Previous work by our laboratory has shown no lasting impact of one-time cervical sterilization on vaginal microbiota [1, 5]. Study timecourse was as follows: pre-IUD phase weeks 0-4, weekly sampling; Cu-IUDs inserted after week 4 sampling; Cu-IUD phase weeks 5-20, sampling every 4 weeks; Cu-IUDs removed after week 20 sampling; post-IUD phase weeks 21-24, weekly sampling. See Fig. S1 for detailed schematic. Baboon perineal staging (from 0-7, with 0-3 being follicular, 4 being ovulation, and 5-7 being luteal, and 7 representing menstruation, with a roughly 30-35 day menstrual cycle) was recorded at each sampling time point, but baboons were not cycle-matched.

### 2.3 Microbiome DNA preparation, sequencing, and analysis

DNA extraction from vaginal swabs was performed with the PowerMag Soil DNA Isolation Kit (Mo Bio, Carlsbad, CA, USA), and amplification of the V4 region of the bacterial 16S rRNA gene, sequencing (Illumina MiSeq), sequence processing using the mothur software package v1.39.5 and analysis were performed as previously described [5]. Sequences were binned into operational taxonomic units (OTUs) based on sequence similarity (3% difference cutoff) using the average neighbor clustering algorithm. Specifically, baboon bacterial communities were compared both between individual animals and within an individual animal over time. Community structure was analyzed using the θ_YC_ dissimilarity coefficient, a measure of both the presence of specific OTU and their relative abundance [6]. Differentially abundant OTUs were identified by LEfSe [7]. Pre-IUD, IUD, and post-IUD timepoints were compared by analysis of molecular variance (AMOVA) [8] on θ_YC_ distances. A mixed effects linear regression model was used to determine the association between perineal staging and θ_YC_.

## 3. Results

### 3.1 Clinical signs

All baboons retained the Cu-IUD for the planned duration (from week 5-20). No animals showed any physical changes or discomfort (as assessed by weight, temperature, pulse, respiration, and frequency of discharge) during the pre-IUD (weeks 0-5), IUD (weeks 6-20), or post-IUD (weeks 21-24) phases of the study, nor did the IUD appear to disrupt the menstrual cycle (data not shown).

### 3.2 Week-to-week vaginal microbial community structure variation

We plotted each animal’s week to week community variability (θ_YC_ distances) with OTUs and their relative abundances at each timepoint, marking menses by coloring the sampling week red (Fig. 1). We compared relative abundances of OTUs within an individual by analysis of molecular variance (AMOVA) [8] and found that animals 3937 and 4051 had no significant differences between pre-IUD, IUD, and post-IUD periods; animal 3903 had a significant difference between the IUD and the pre-IUD periods (p = 0.035); animals 3913 and 4047 had significant differences between the post-IUD and pre-IUD periods (p = 0.032 and 0.025, respectively); and animal 3913 had a significant difference between the post-IUD and IUD periods (p = 0.012).

**Figure 1.**
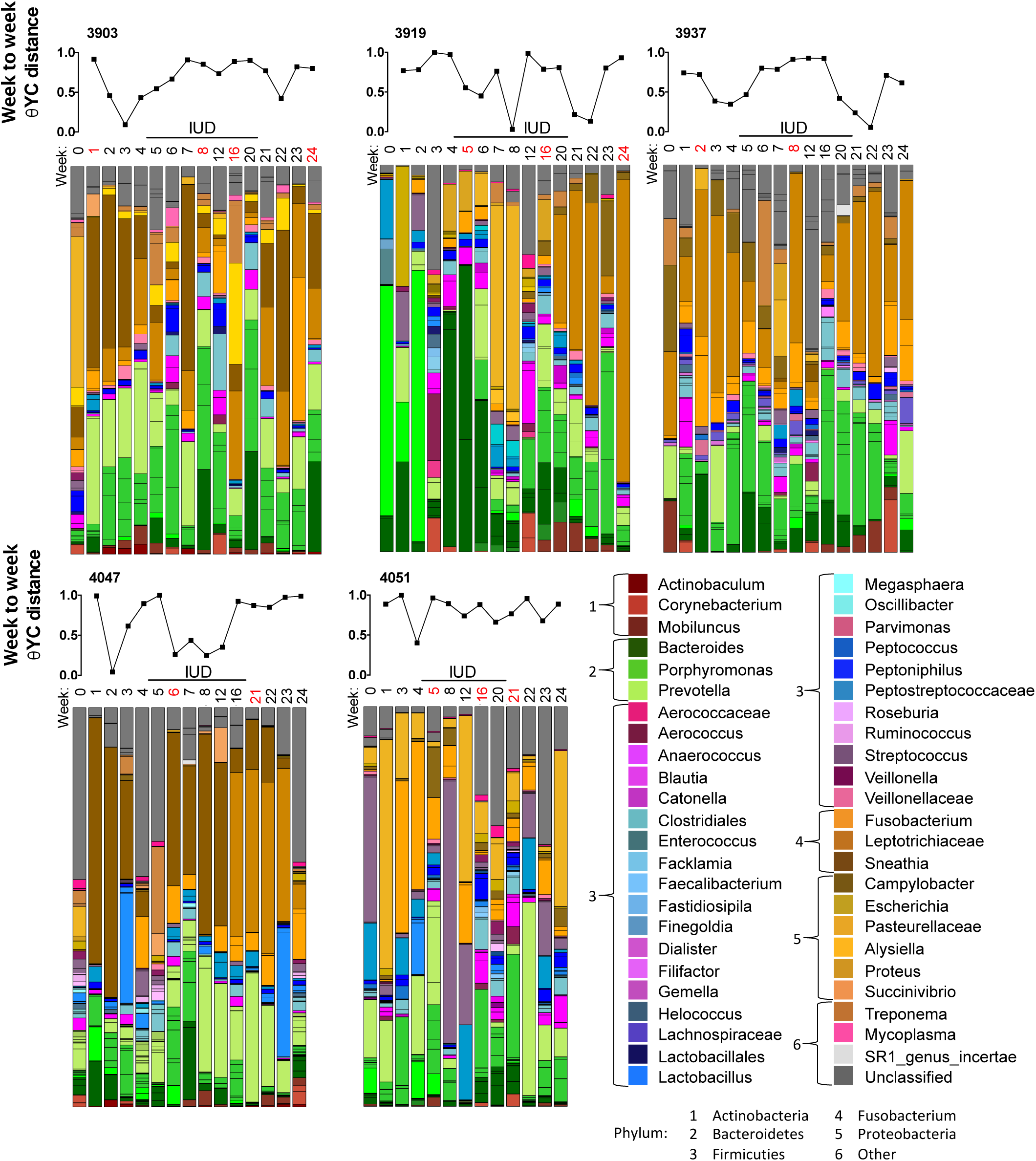
Bacterial communities exhibit inter-animal variation, but no pattern in change between pre-IUD, IUD, and post-IUD periods. Week-to-week change in θYC values in each animal is matched to the bar chart of microbial communities at each timepoint. Microbial communities are color coded as indicated, and are grouped by phylum indicated. LEfSe analysis performed on each individual animal by grouping communities found in the pre-IUD, IUD, and post-IUD timepoints found: animal 3903 had significant difference in relative abundance between the pre-IUD and IUD phases; animal 3919 had a significant difference in relative abundance between the IUD and post-IUD timepoints; animals 3919 and 4047 had shifts in relative abundance of microbial communities between the pre-IUD and post-IUD periods. Menses are indicated by red week numbers.

Examination of the average θ_YC_ distance showed high within-animal week-to-week variation in community structure; community stability, measured by week-to-week θ_YC_ distances, did not change significantly between pre-IUD, IUD, and post-IUD timepoints when taken as a group (Fig. S2), which was independent of an individual animal’s menstrual staging (p = .1441, Fig. S3). The greatest shift in bacterial community composition, indicated by the highest θ_YC_ distances, occurred between week 0 and week 1, when the animals were moved from group housing to single cages (Fig. S2).

### 3.3 Microbial community composition and clustering

The most frequent bacterial phyla as grouped and ranked by mothur across all baboons were Bacteroidetes, Fusobacteria, Proteobacteria, and Firmicutes (Fig. 1). Comparing the pre-IUD, IUD, and post-IUD timepoints of all animals by AMOVA on θ_YC_ distances, we found no significant differences due to insertion or long-term use of the IUD (p-value: 0.126); however, there was a significant difference between post-IUD communities and both pre-IUD communities (p-value: 0.003) and IUD communities (p-value: <0.001) (Fig. 2).

**Figure 2.**
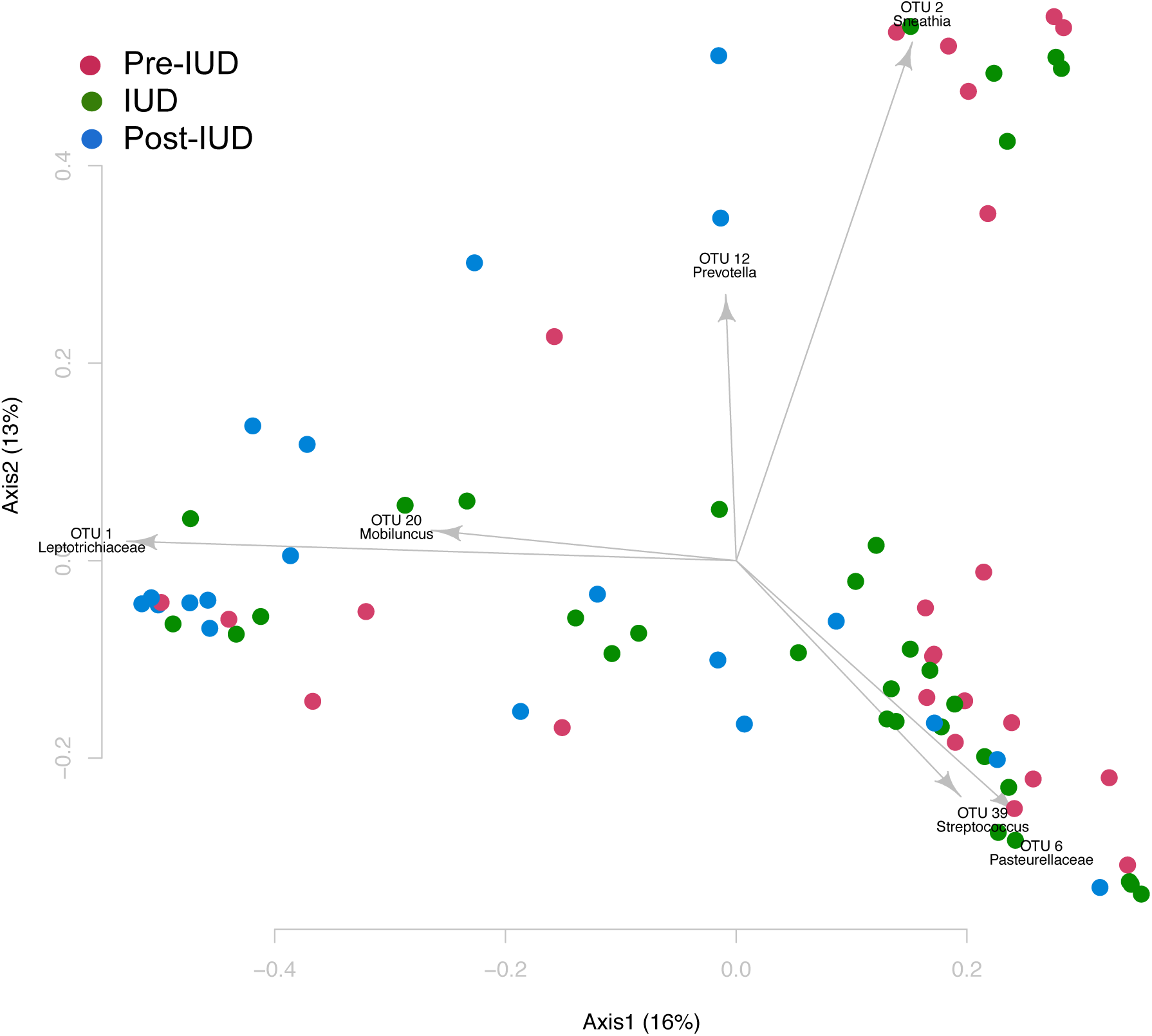
Principal Coordinates Analysis (PCoA) shows no distinct clustering of OTUs with regard to IUD insertion but changes after IUD removal. Baboons at each timepoint are represented by a circle, colored to indicate pre-IUD, IUD, and post-IUD periods. Axes indicate percentage of variation by the plotted principal coordinates. Operational Taxonomic Units (OTUs) of related bacteria are overlaid on the data. Significant differences (in OTU1 Leptotrichaceae) were seen by AMOVA when post-IUD timepoints were compared to both pre-IUD and IUD timepoints.

Differentially abundant OTUs included OTU1 (Leptotrichiaceae), which was higher in the post-IUD timepoints relative to both pre-IUD and IUD (Fig. S4, see table Table S1 for OTU identities of Fig. 2, S4). When we followed community structure over time, communities within an individual were more similar then those found in different animals (Fig. S5, Table S1 for OTU identities).

## 4. Discussion

This study was designed to assess whether transcervical insertion of Cu-IUD was possible, measure any clinical changes due to Cu-IUD placement in baboons, and measure changes in the vaginal microbiota due to Cu-IUD placement, use, or removal. We found no clinical changes to baboons (*e.g*. temperature, weight, menstrual cycle, discharge) and no changes in the vaginal microbiota attributable to Cu-IUD insertion and long-term use; however, we did find a slight but statistically significant alteration of the vaginal microbiota after Cu-IUD removal. The baboon recapitulates data and reports from human patients receiving a Cu-IUD: transcervical insertion is successful, no physiologic changes are observed while using the Cu-IUD, and insertion and use do not significantly affect the vaginal microbiota [9]. As we have demonstrated with the LNG-IUS in the baboon, we conclude that the olive baboon is a feasible model system for Cu-IUD studies.

The lack of change between the pre-IUD phase and IUD phase is consistent with recently published work in humans, although the human study did not extend to after IUD removal [9]. The mechanism behind the change in vaginal microbiota after IUD removal is unclear, as is the biological significance. Two baboons (3913 and 4047) are largely the driving force behind post-IUD removal vaginal microbiota changes, while the other 3 showed no significant changes associated with removal.

Animal models of Cu-IUD are limited. In mice, insertion of a copper wire into the uterine horn [10] is limited by the reliance on surgery, the vastly different estrus cycle and reproductive tract physiology from humans, and the more limited range of naturally-occurring sexually-transmitted pathogens relative to humans and NHP. NHP studies of Cu-IUD have been successfully performed in the pigtail macaque [11], but surgery is necessary for device insertion, which carries significant risks to the animal. Thus, the primary strength of this study is the development and characterization of an animal model for studying Cu-IUD in a large NHP whose reproductive tract is analogous to the human reproductive tract– including an easily monitored 30-35 day menstrual cycle, and a straight cervical canal– and which has no alterations in insertion from humans. There are several limitations of this study. Comparisons between animals are limited by the small sample size and the lack of cycle matching. Cycle matching animals for IUD insertion would eliminate one variable in changing vaginal microbial communities, but due to the idiosyncratic nature of vaginal microbiota, this is not guaranteed to result in reduced variability between baboon theta-YC distances. There is no clear association in these animals between cycle stage and theta-YC distance. Additionally, the study ended 4 weeks into the post-IUD removal period, which we surprisingly found to be the only major shift in microbial community structure. Future studies investigating microbial community shifts post-IUD removal should extend the post-IUD removal sampling period to determine if vaginal microbial communities return to baseline and if so, how long this takes.

The diverse vaginal microbiota in baboons and the common lack of Lactobacillus dominance is reminiscent of bacterial vaginosis in female patients. One limitation to the use of baboons for bacterial vaginosis studies is that several of the clinical (Amsel) criteria for diagnosis of bacterial vaginosis– odor and discharge– are difficult to assess from a baboon; however, criteria based on bacterial species present on gram stain (Nugent scores) or sequencing data are easily obtained.

Development of this baboon model of Cu-IUDs opens many avenues of important investigation which are ethically impossible in humans: 1) the interaction of Cu-IUDs with HIV, for which baboons are a viable model [12]; 2) investigation of the often-reported association between Cu-IUDs and bacterial vaginosis, as the high pH and diverse microbial community of the baboon mimic aspects seen in human bacterial vaginosis [13]; and 3) the impact of Cu-IUDs on endometriosis [14]. Our study opens a new avenue of research into the intersection of the Cu-IUD and human reproductive tract disorders and pathogens.

## 5. Acknowledgements

The authors would like to acknowledge the Human Microbiome Initiative.

## Supplemental Figure Legends

**Figure S1.**
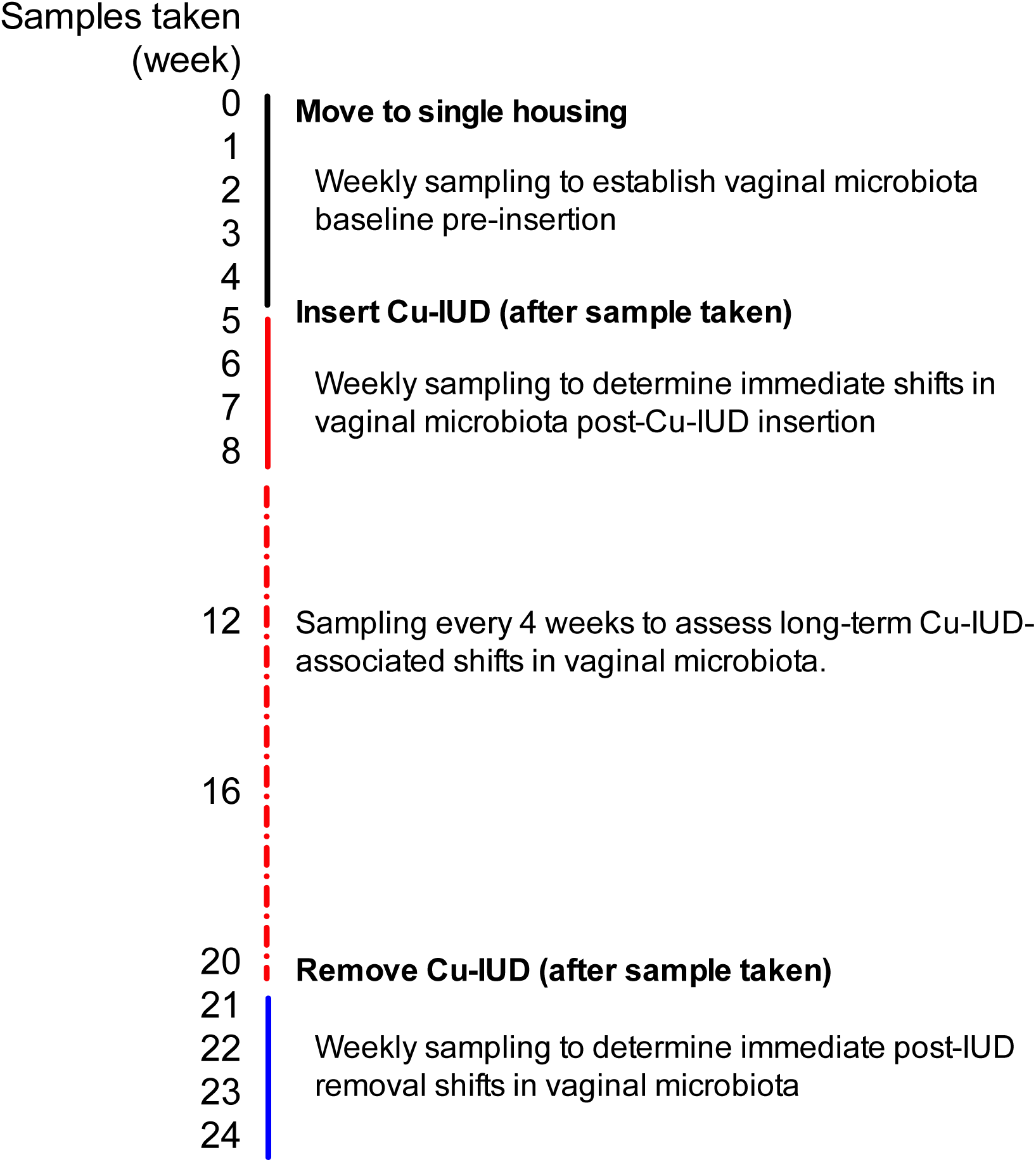
Sampling schematic.

**Figure S2.**
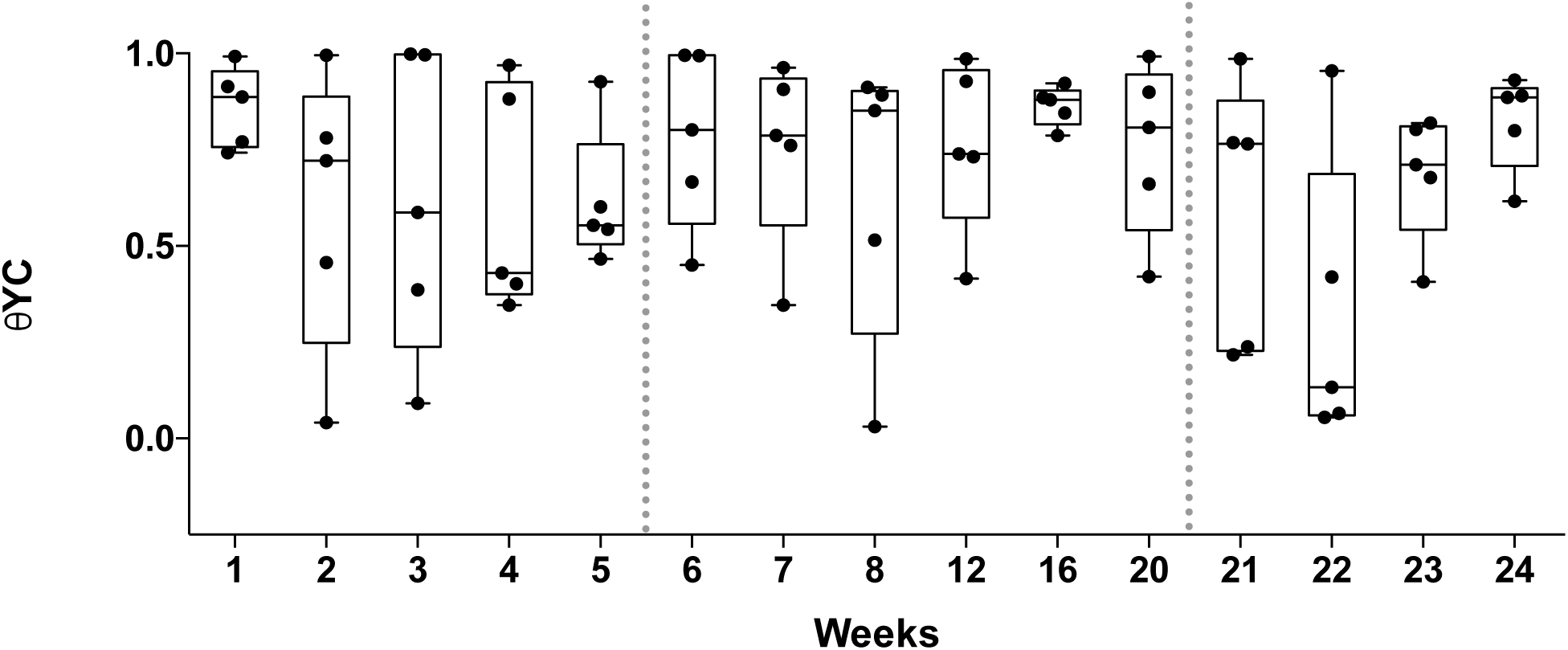
Averaged week-to-week θYC values do not change significantly between pre-IUD, IUD, and post-IUD periods.

**Figure S3.**
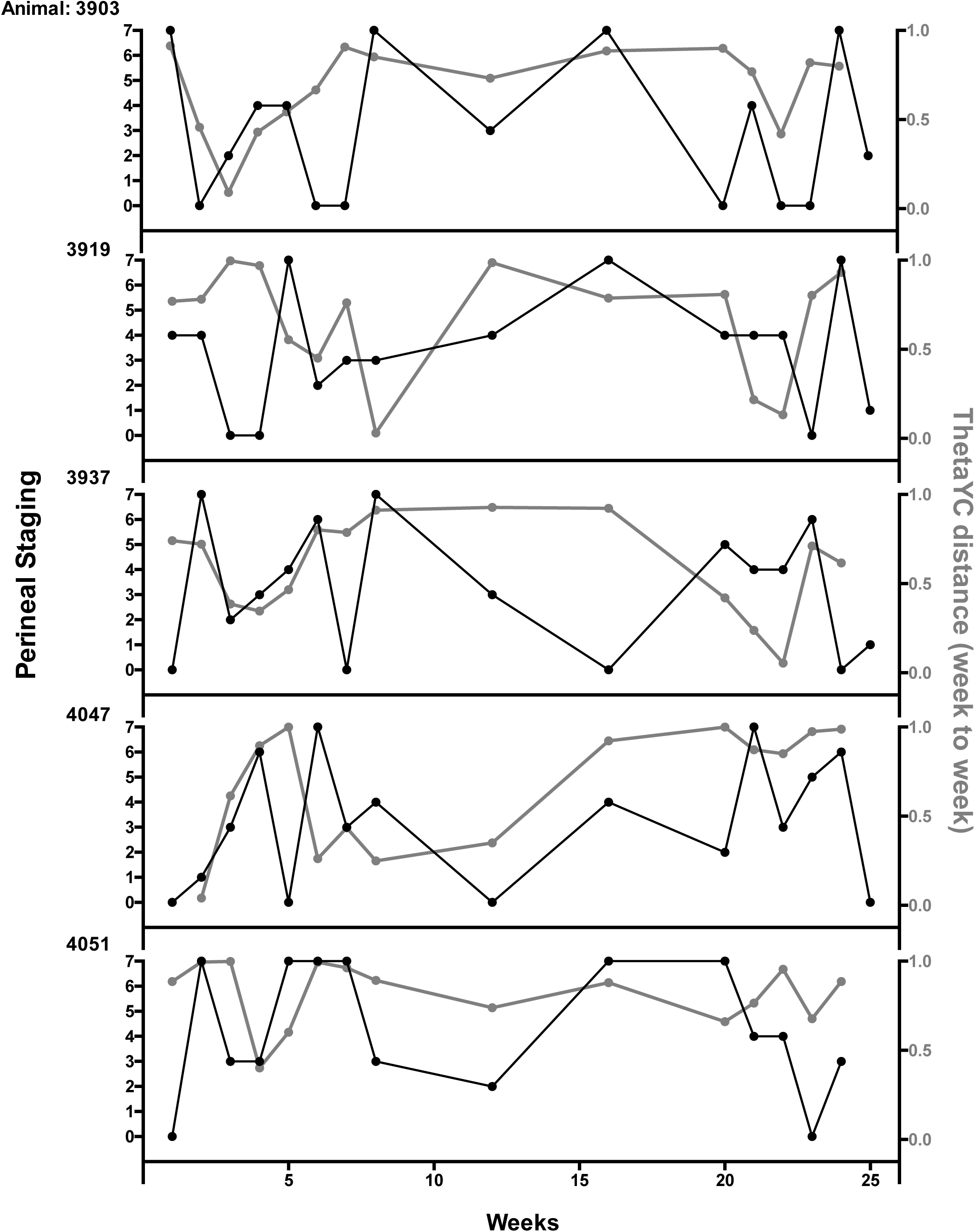
Week-to-week θYC distances are independent of baboon perineal stagimg. Perineal staging of baboons by the Institute for Primate Research stages baboon menstrual cycle from 0-7, with 0-3 being follicular, 4 being ovulation, and 5-7 being luteal, and 7 representing menstruation. Animals were not stage-matched. We performed mixed effects modeling and multivariate analysis of our data matching perineal staging to θ_YC_ thetaYC distances and found no significant association using either method.

**Figure S4.**
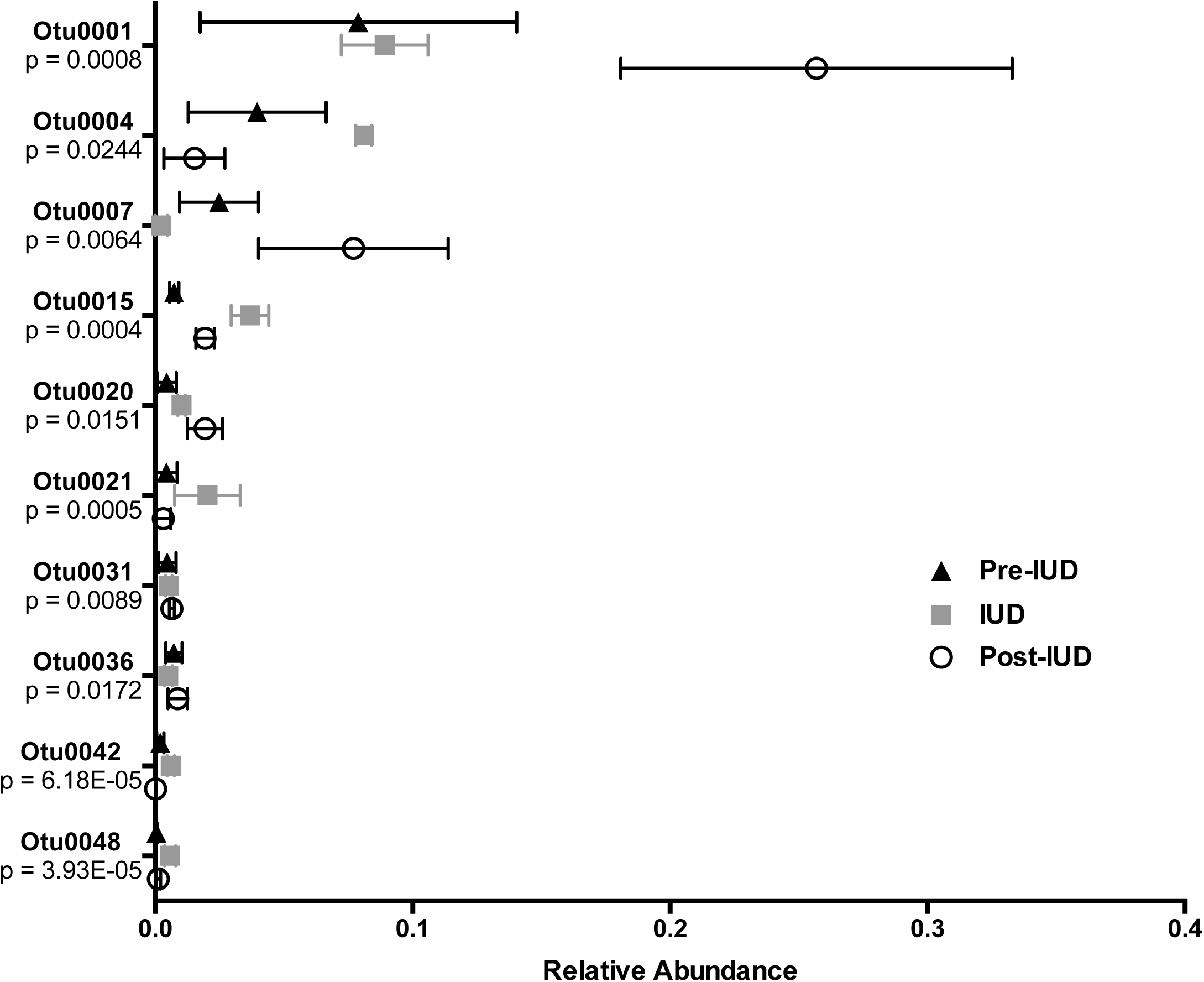
Significant differences by LEfSe in 10 OTUs with abundance greater than 0.001 in any group. OTU number and p-value (by LEfSe) are indicated on the Y-axis and the relative abundance is indicated on the X-axis. LEfSe analysis was a 3-way comparison between timepoints pre-IUD, IUD, and post-IUD. When further analyzed by 2-way ANOVA with multiple comparisons: OTU0001 was significantly different from pre-IUD and IUD; OTU0004 was significantly different in post-IUD relative to IUD; and OTU0007 was significantly different in post-IUD relative to IUD. Our cutoff for inclusion was OTUs with a relative abundance greater than 0.001 in any group.

**Figure S5.**
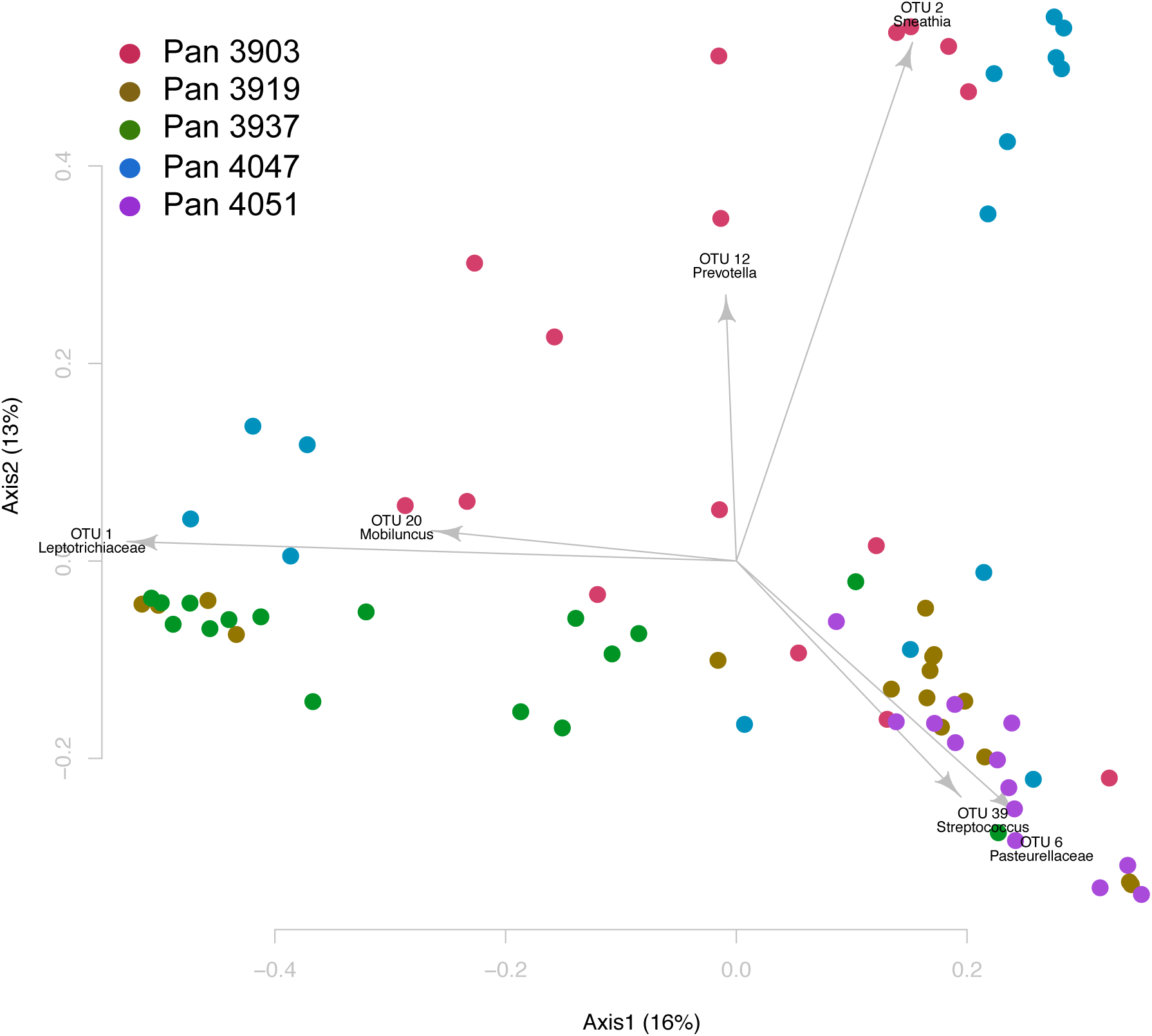
Principal Coordinates Analysis (PCoA) show animals are most similar to themselves and change little over 24 weeks. Baboons at each timepoint are represented by a circle, colored to indicate individual animals. Axes indicate percentage of variation by the plotted principal coordinates. Operational Taxonomic Units (OTUs) of related bacteria are overlaid on the data. No significant differences were seen by AMOVA.

**Table S1.**
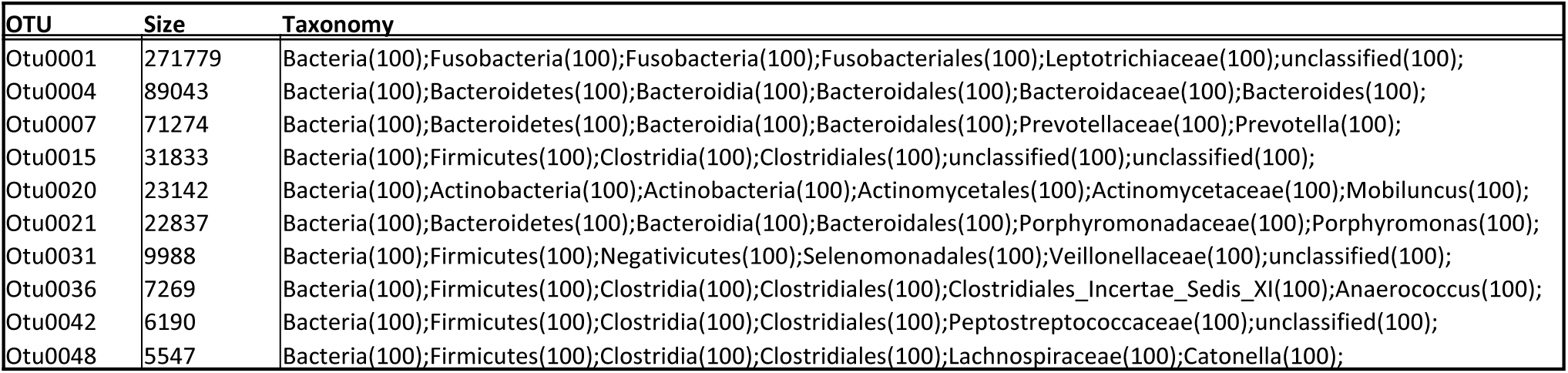
OTU identities from Fig. 2, Fig. S2, and Fig. S5 for groups with significant differences in abundance between timepoints determined by LEfSE.

## References

[1] Bell JD, Bergin IL, Natavio MF, et al. Feasibility of LNG-IUS in a baboon model. Contraception. 2013;87:380–4.

[2] Madden T, Grentzer JM, Secura GM, Allsworth JE, Peipert JF. Risk of bacterial vaginosis in users of the intrauterine device: a longitudinal study. Sexually transmitted diseases. 2012;39:217–22.

[3] Hashway S. Impact of a Hormone-Releasing Intrauterine System on the Vaginal Microbiome: A Prospective Baboon Model. Journal of medical primatology. 2013:89–99.

[4] Bauer C. The baboon (Papio sp.) as a model for female reproduction studies. Contraception. 2015;92:120–3.

[5] Liechty ER, Bergin IL, Bassis CM, et al. The levonorgestrel-releasing intrauterine system is associated with delayed endocervical clearance of Chlamydia trachomatis without alterations in vaginal microbiota. Pathogens and disease. 2015;73:ftv070.

[6] Yue JC, Clayton MK. A Similarity Measure Based on Species Proportions. Communications in Statistics - Theory and Methods. 2005;34:2123–31.

[7] Segata N, Izard J, Waldron L, et al. Metagenomic biomarker discovery and explanation. Genome biology. 2011;12:R60.

[8] Anderson MJ. A new method for non-parametric multivariate analysis of variance. Austral Ecology. 2001;26:32–46.

[9] Bassis CM, Allsworth JE, Wahl HN, Sack DE, Young VB, Bell JD. Effects of intrauterine contraception on the vaginal microbiota. Contraception. 2017;96:189–95.

[10] Chou CH, Chen SU, Shun CT, Tsao PN, Yang YS, Yang JH. Divergent endometrial inflammatory cytokine expression at peri-implantation period and after the stimulation by copper intrauterine device. Scientific reports. 2015;5:15157.

[11] Engel RM, Morris M, Henning T, et al. Evaluation of pigtail macaques as a model for the effects of copper intrauterine devices on HIV infection. Journal of medical primatology. 2014;43:349–59.

[12] Locher CP, Witt SA, Herndier BG, Tenner-Racz K, Racz P, Levy JA. Baboons as an animal model for human immunodeficiency virus pathogenesis and vaccine development. Immunological reviews. 2001;183:127–40.

[13] Rivera AJ, Frank JA, Stumpf R, et al. Differences between the normal vaginal bacterial community of baboons and that of humans. American Journal of Primatology. 2011;73:119–26.

[14] Braundmeier AG, Fazleabas AT. The non-human primate model of endometriosis: research and implications for fecundity. Molecular human reproduction. 2009;15:577–86.

